# Immune gene diversity in archaic and present-day humans

**DOI:** 10.1101/348581

**Authors:** David Reher, Felix M. Key, Aida M. Andrés, Janet Kelso

## Abstract

Genome-wide analyses of two Neandertals and a Denisovan have shown that these archaic humans had lower genetic heterozygosity than present-day people. A similar reduction in genetic diversity of protein-coding genes (*gene diversity*) was found in exome sequences of three Neandertals. Reduced gene diversity, and particularly in genes involved in immunity, may have important functional consequences. In fact, it has been suggested that reduced diversity in immune genes may have contributed to Neandertal extinction. We therefore explored gene diversity in different human groups and at different time points on the Neandertal lineage with a particular focus on the diversity of genes involved in innate immunity and genes of the Major Histocompatibility Complex (MHC).

We find that the two Neandertals and the Denisovan have similar gene diversity, both significantly lower than any present-day human. This is true across gene categories, with no gene set showing an excess decrease in diversity compared to the genome-wide average. Innate immune-related genes show a similar reduction in diversity to other genes, both in present-day and archaic humans. There is also no observable decrease in gene diversity over time in Neandertals, suggesting that there may have been no ongoing reduction in gene diversity in later Neandertals, although this needs confirmation with a larger sample size. In both archaic and present-day humans, genes with the highest levels of diversity are enriched for MHC-related functions. In fact, in archaic humans the MHC genes show evidence of having retained more diversity than genes involved only in the innate immune system.

## Introduction

Since the first complete Neandertal genome was sequenced (Green et al. 2010), ongoing efforts have retrieved DNA sequences from a number of additional extinct hominins (Castellano et al. 2014; Meyer et al. 2012; Prufer et al. 2017; Prufer et al. 2014). Comparing the Neandertal and Denisovan genomes to the genomes of present-day people provided evidence that the ancestors of all non-Africans living today met and interbred with Neandertals (Green et al. 2010) and that the ancestors of people that today live in Oceania interbred with Denisovans (Reich et al. 2010). Although some of the resulting introgressed DNA has been shown to be adaptive in anatomically modern humans (Dannemann et al. 2016; Huerta-Sanchez et al. 2014; Racimo et al. 2017; Racimo et al. 2015), conserved regions of present-day human genomes are significantly depleted of introgressed Neandertal sequence, which has been interpreted as evidence for purifying selection against introgressed Neandertal DNA in anatomically modern human genomes (Fu et al. 2016; Harris and Nielsen 2016; Juric et al. 2016; Sankararaman et al. 2014). Recent studies suggested that slightly deleterious alleles may have accumulated in the genomes of Neandertals and Denisovans because of reduced efficacy of natural selection as a result of their small long-term effective population size (N_e_) (Harris and Nielsen 2016; Juric et al. 2016).

All archaic individuals analysed to date have genome-wide heterozygosities that are lower than those seen in present-day humans. The genome-wide heterozygosity of a ~50,000-year-old Neandertal from Vindija cave in Croatia (Prufer et al. 2017) was estimated to be 1.6×10^−5^, similar to that previously reported for the ~120,000-year-old Altai Neandertal (Prufer et al. 2014) and only slightly lower than the estimate for an ~80,000-year-old Denisovan individual (1.8 x10^−5^) (Meyer et al. 2012). This low genetic diversity has also been observed in the exome sequences of three Neandertals from the Vindija, El Sidrón and Denisova Caves which show lower average heterozygosities than present-day humans (Castellano et al. 2014). Genetic diversity in genic regions is particularly important, as it can potentially impact the levels of functional diversity in the population. However, the limited number of high quality archaic genome sequences means that we do not know to what extent levels of gene diversity (i.e., genetic diversity in protein-coding genes) may have changed over time. The availability of two high-coverage Neandertal genomes of individuals who lived 70,000 years apart, as well the high-coverage genome of one Denisovan, now allow us to begin to explore gene diversity in archaic human populations at different times during Neandertal history.

It has been suggested that lack of functional variation in immune-related genes – especially in genes related to the innate immune system which is known to serve as a first defence mechanism against pathogen detection –, some of which are targets of long-term balancing selection (Bitarello et al. 2018; Key et al. 2014b; Meyer and Thomson 2001), could have contributed to Neandertal extinction (Houldcroft and Underdown 2016; Sullivan et al. 2017; Wolff and Greenwood 2010). This is known as the differential pathogen resistance hypothesis (DPRH), and (Sullivan et al. 2017) recently presented evidence both for and against this hypothesis. Using the exome data (Castellano et al. 2014), they found that Neandertals had substantially lower numbers of non-synonymous single nucleotide polymorphisms (SNPs) than present-day humans in 73 innate immune-related genes, 12 genes of the Major Histocompatibility Complex (MHC), 164 virus-interacting protein genes, and 73 loci with high diversity in chimpanzee (which might be enriched for targets of balancing selection). They concluded that reduced protein sequence diversity in this set of immune genes may have resulted in reduced resistance to pathogens and have thereby contributed to Neandertal extinction. However, on the other hand they also reported a higher number of non-synonymous SNPs in Neandertals than in present-day humans for 12 genes of the MHC, suggesting high levels of functional diversity in this component of the immune system.

Here, we leverage existing high-quality whole-genome data from three archaic humans to test for evidence of a specific reduction of gene diversity in archaic humans that would be expected under the DPRH. We focus on comparing genetic diversity between archaic and present-day humans, and over time in the Neandertal lineage in a comprehensive set of 1,548 innate immunity genes. Genes of the innate immune system and their evolutionary history have been well characterized and include a rather well-defined set of genes that are not affected by infection history (Deschamps et al. 2016; Quintana-Murci and Clark 2013). In addition, we include 14 MHC genes because of their important role in immunity, and their well-studied and unique evolutionary history (Bitarello et al. 2018; Key et al. 2014b; Meyer and Thomson 2001). In a second analysis, we generalise the idea underlying the DPRH. Instead of exploring gene diversity only in innate immunity genes, we tested if any functional category of genes had particularly high or low gene diversity in Neandertals when compared with modern humans.

## Methods

### Data

Our analyses are based on three published high-coverage genomes of the Altai and Vindija Neandertals and the Denisovan (Meyer et al. 2012; Prufer et al. 2017; Prufer et al. 2014) as well as a published data set of 14 present-day individuals consisting of five individuals from Africa (Mandenka, Mbuti, San, Yoruba, Dinka), three from Asia (Dai, Han, Papuan), two from Australia, two from Europe (French, Sardinian) and two from South America (Karitiana, Mixe) (Meyer et al. 2012). For all analyses, we used the filters applied by (Prufer et al. 2017). In brief, we retained sequences with mapping quality higher than 25, sites with coverage higher than 10 (including both a 2.5% higher and lower coverage cut-off; corrected for GC content), and unique positions in the genome according to 35-mer 1-mismatch filter, while removing simple repeats (tandem repeat finder track at UCSC). We downloaded a list of all annotated autosomal human protein-coding genes from BioMart/Ensembl Release 84 (GRCh37) (Yates et al. 2016) including introns, exons, and additional 1kb up and downstream to capture adjacent regulatory elements, and filtered for uniqueness by HGNC symbol and gene coordinates (GRCh37/hg19, N = 17,505). We extracted the sequences that pass these filters for each individual from the whole genome VCF files and excluded genes with fewer than 2,000 callable sites from the analysis (reducing the number of genes by 2,041 genes for each individual on average). Data processing was done using Tabix (Li 2011), BEDOPS (Neph et al. 2012) and BEDTools (Quinlan 2014) and statistical analysis and visualisation was done using R (Team 2016).

### Measure of gene diversity

To estimate gene diversity for autosomal protein-coding genes per genome, we counted single nucleotide polymorphisms (SNPs) (Andres et al. 2009). For individual genomes, a SNP is defined as a biallelic heterozygous site. To account for local heterogeneity in mutation rate and the rate of substitutions, we divided the number of SNPs by the number of fixed differences (FDs) which served as a proxy for mutation rate. For individual genomes, a FD is a site in which the chimpanzee reference allele (taken from the EPO alignment version 69 (Yates et al. 2016) based on the chimpanzee reference CHIMP2.1.4) is different from a homozygous allele in the test individual. We calculated the SNP/FD ratio for each gene that passed our filter and define this measure as a proxy for genetic diversity in protein-coding genes which we call gene diversity. We note that because we consider the full length of genes including regulatory sequences and introns, and use divergence with chimpanzee, the number of genes with very small number of FDs is extremely low (on average, there are eight genes with less than three FDs per individual; the mean number of FDs per gene over all individuals is 272). Further, we pre-defined sets of genes (more specifically, innate immune and MHC genes, see below), summed the total number of SNPs in those genes, and divided that number by the total number of FDs in those genes to a combined single ratio of SNPs to FDs per individual (mean SNP/FD ratio). We computed confidence intervals on bootstrapped sets. After sampling N genes with replacement from both test and background gene sets, we re-calculated the SNP/FD ratios for each of 5,000 resampled sets and defined cut-offs based one the 2.5% and 97.5% quantiles of the resulting empirical distributions as cut-offs.

### Diversity in innate immune and MHC genes

To test whether there is evidence for an overall increase or reduction in gene diversity of innate immune genes in archaic humans, we calculated SNP/FD ratios in a comprehensive set of innate-immune genes curated by (Deschamps et al. 2016). This set combines genes from InnateDB (Breuer et al. 2013) and genes assigned to the GO category *innate immune response* (GO:0045087). We updated this gene list with recent InnateDB entries following our filtering scheme. This resulted in a set of 1,548 innate immunity genes. We additionally investigated the following subsets of this innate-immune gene list separately: *toll-like receptor signalling pathway* (GO:0002224, N = 169), *innate immune response in mucosa* (GO:0002227, N = 10), *defense response* (GO:0006952, N = 65), *defense response to bacterium* (GO:0042742, N = 60), *defense response to Gram-negative bacterium* (GO:0050829, N = 29), *defense response to Gram-positive bacterium* (GO:00050830, N = 52), *defense response to fungus* (GO:0050832, N = 16), *defense response to virus* (GO:0051607, N = 154), as well as the MHC genes (N = 14). We then defined a hand-curated set of autosomal protein-coding background genes without any reported immune function to use as a background set (13,393 Ensembl genes (Yates et al. 2016)) for which we excluded 4,723 genes with any reported immune system-related function (ImmPort gene list (Breuer et al. 2013)) as well as genes shorter than 500 bp in length. To compare the diversity of immune genes relative to the protein-coding background between archaic and present-day humans, we normalised the levels of gene diversity for each individual by the overall gene diversity in the set of background genes found in that same individual, i.e. we divided the mean SNP/FD ratio of innate-immune genes by the mean SNP/FD ratio of the background genes (normalised gene diversity). We repeated the same analysis for the MHC gene set, i.e. all *HLA* genes on chromosome 6.

### GO enrichment analysis

We performed a gene ontology enrichment analysis to explore whether any particular functional groups of genes (Gene Ontology categories, GO) are overrepresented among the genes with the highest (top-5% tail of the empirical SNP/FD ratio distribution) or lowest (bottom-5% tail of the same distribution) SNP/FD ratios in the three archaics, or a set of three representative present-day humans (Africa (Yoruba), Europe (French) and Asia (Han)). In this analysis, we only considered genes that pass our above-mentioned filters in the test individuals, and averaged the SNP/FD ratio over those individuals for each gene. For GO enrichment analyses, we used the R package ‘GOfuncR’ (Grote 2017; Grote et al. 2016; Prufer et al. 2007). In the GO enrichment analyses, we compared the test sets to all genes with SNP/FDs ratios outside the top and bottom-5% in the relevant set of three individuals. We further performed GO enrichment analyses for pairs of genomes and for individual genomes. While these analyses have lower power than the one above, they allow us to better define and understand the enrichment signal in genes with specific functions.

## Results

### Archaic humans had lower overall gene diversity than present-day humans

We estimated gene diversity per individual by calculating SNP/FD ratios in five present-day individuals from African populations (Mandenka, Mbuti, San, Yoruba and Dinka), and nine present-day individuals from non-African populations (French, Sardinian, Dai, Han, Papuan, Karitiana, Mixe, and two Australians). In agreement with previous observations, we consistently found significantly higher diversity in African individuals than in individuals from non-African populations (Figure 1 (a), indicated by non-overlapping 95% confidence intervals), consistent with reduced diversity in non-Africans as a consequence of the out-of-Africa bottleneck (reviewed in (Cavalli-Sforza and Feldman 2003)). All three archaic humans exhibit significantly lower gene diversity compared to the present-day humans, consistent with their previously reported overall low genomic diversity (Castellano et al. 2014; Meyer et al. 2012; Prufer et al. 2017; Prufer et al. 2014). The Altai Neandertal has lower gene diversity than the other two archaic individuals likely as a consequence of recent inbreeding (Prufer et al. 2014). Removing the extended tracts of homozygosity (defined by (Prufer et al. 2017; Prufer et al. 2014)) from the Altai genome (Altai*), results in comparable levels of gene diversity in the Altai and Vindija Neandertals. Gene diversity in both of the Neandertals is slightly lower than in the Denisovan which is consistent with the reported differences in genome-wide diversity (Meyer et al. 2012). [Approximate position of Figure 1]

**Fig. 1:**
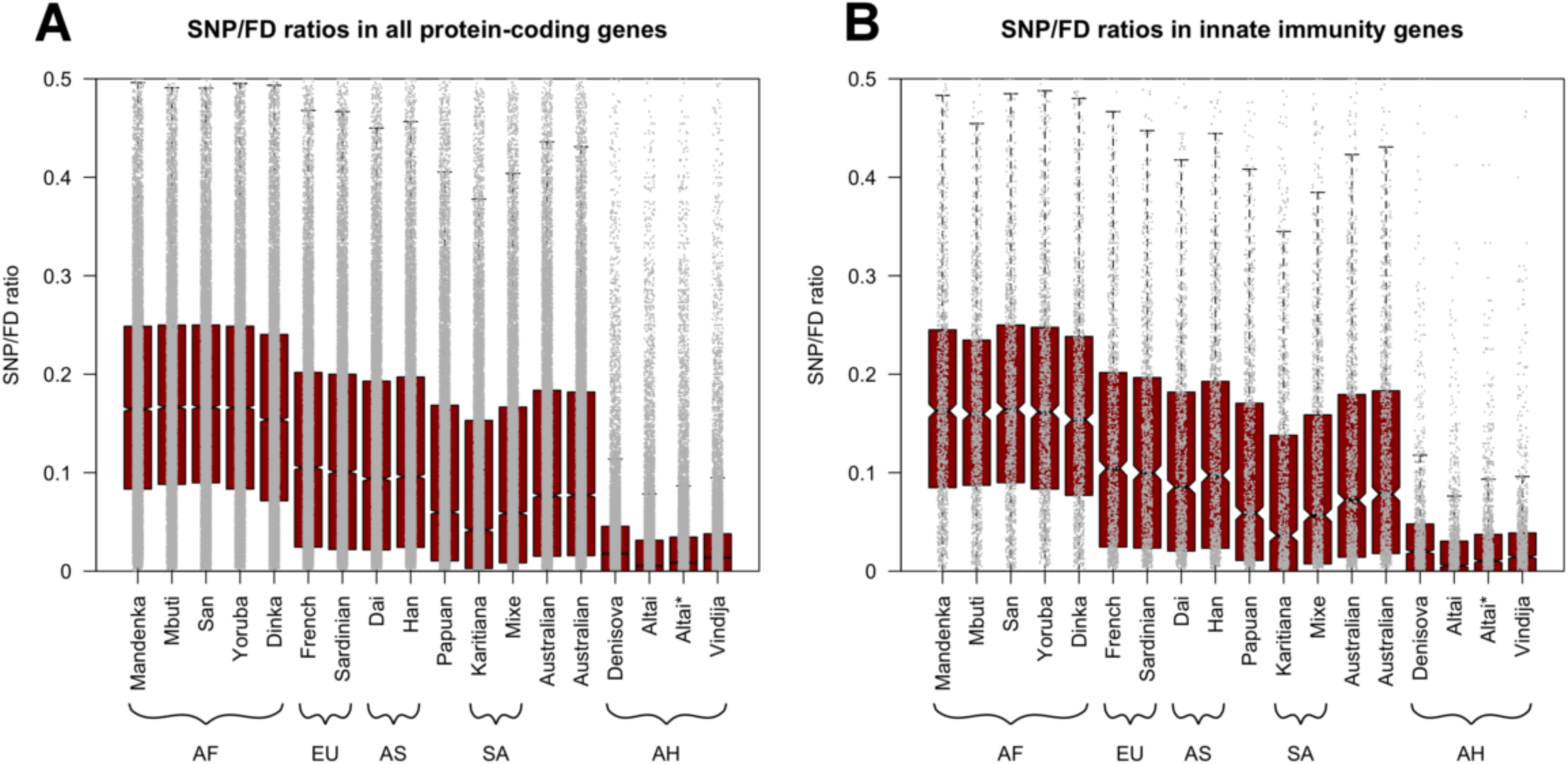
Distributions of SNP/FD ratios per gene for all 17 individuals. Black lines and notches give medians and 95% confidence intervals, respectively. Y-axis trimmed at 0.5 for clarity (for full plots see Figure S1), each grey dot gives the SNP/FD ratio for a single gene. **(A)** All protein-coding genes and **(B)** innate immune-related genes. AF = African, EU = European, AS = Asian, SA = South American, AH = Archaic Human.

### Archaic humans had similarly low gene diversity in innate immune genes compared to non-immune genes

Next, we tested whether genes of the innate immune system showed a similar reduction in Neandertals (when compared with present-day humans) as non-immune genes. Figure 1 (b) shows the distribution of SNP/FD ratios for each immune gene in all individuals as well as for the Altai Neandertal excluding homozygous tracts (Altai*). As was the case for all autosomal protein-coding genes (Figure 1 (a)), present-day humans from Africa (Mandenka, Mbuti, San, Yoruba and Dinka) have higher diversity in innate immune-related genes (SNP/FD ratios range from 0.154 to 0.167) than individuals from non-African populations (French, Sardinian, Dai, Han, Papuan, Karitiana, Mixe, and two Australians, SNP/FD ratios range from 0.042 to 0.105). With values from 0.005 to 0.018, the median SNP/FD ratios are lower for the three archaic humans than for the present-day humans (Figure 1 (b)). The median SNP/FD ratio for the Altai Neandertal is slightly (not significantly) lower than that for the Vindija 33.19 Neandertal. Again, after removing identified homozygous tracts, the Altai Neandertal (Altai*) exhibits similar gene diversity to the younger Vindija 33.19 Neandertal and the Denisovan, suggesting the lower SNP/FD ratio is likely a result of recent inbreeding in the Altai Neandertal (Prufer et al. 2014).

To further investigate a putative specific reduction in innate immune gene diversity, we investigated normalised immune gene diversity by dividing mean SNP/FD ratios in immune genes by mean SNPD/FD ratios in a set of non-immune-related background genes (see Methods, Figure 2). There are no significant differences in the normalised gene diversities between any pair of ancient or present-day individuals, as indicated by overlapping 95% confidence intervals. Furthermore, 95% confidence intervals include the value 0 in 16 of the 17 individuals (in one Australian the upper confidence interval limit is slightly below 0). This suggests that in all individuals, innate-immune related genes have levels of diversity that are expected given their genome-wide gene diversity. We find thus no indication that innate immunity genes in archaic individuals have significantly different levels of normalised gene diversity than in present-day humans. These results are also reflected in the analysis of eight subsets of innate immunity genes (containing 10 to 169 genes, respectively) in which we also find no evidence for a specific reduction of gene diversity, even though there is some variation due to low sample size (Figure S2).

**Fig. 2:**
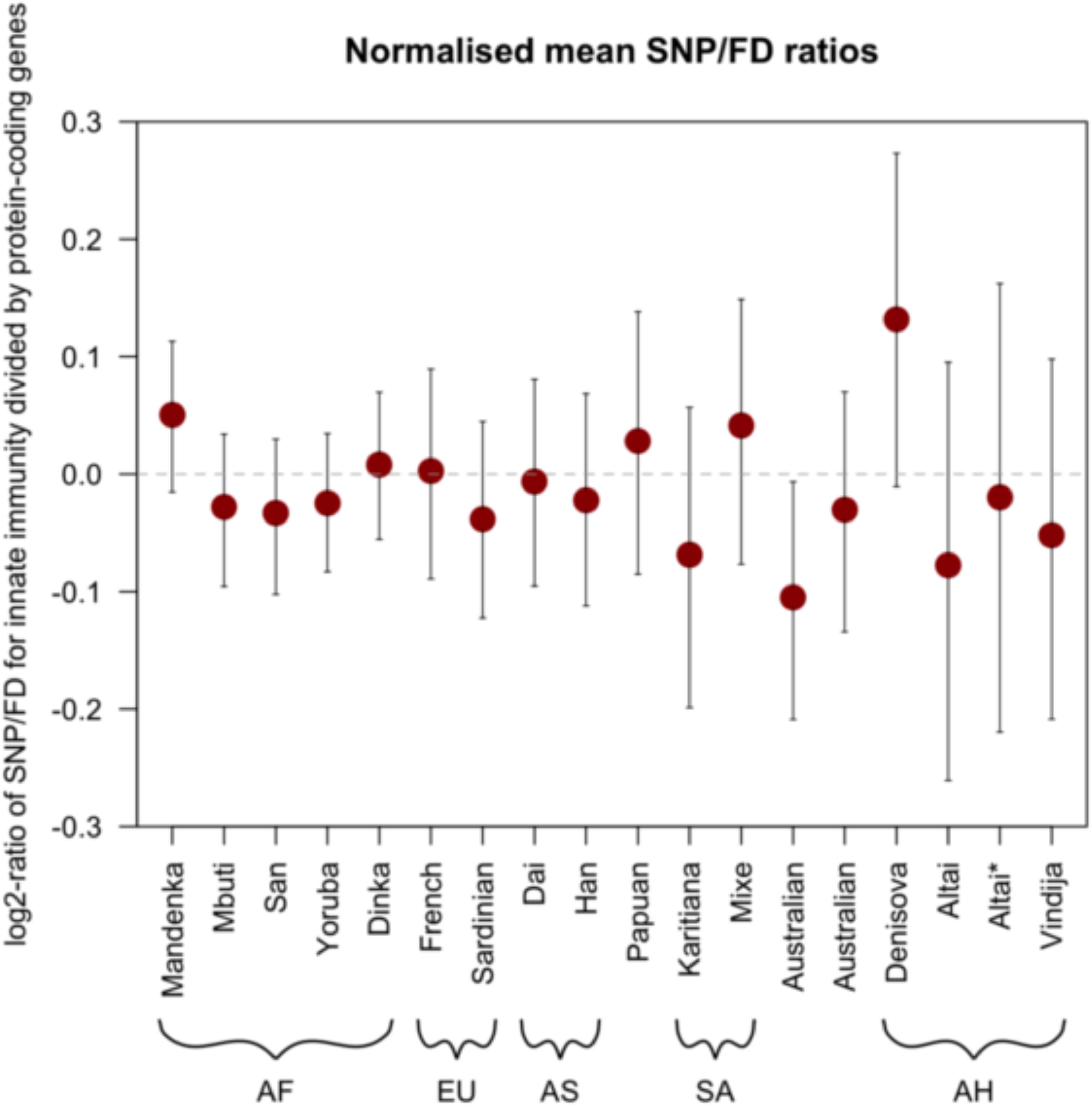
Normalised mean SNP/FD ratios (log2) for all 17 individuals in the full set of innate immune-related genes (N = 1,548). Error bars give 95% confidence intervals calculated by bootstrapping (B = 5,000). AF = Africa, EU = European, AS = Asian, SA = South American, AH = Archaic Human. Dashed line gives expected value if the mean values for innate immunity genes and autosomal protein-coding background genes were equal.

The younger Vindija 33.19 Neandertal and the older Altai Neandertal show similar levels of normalised gene diversity in innate immune genes (−0.05 and −0.02 for Vindija 33.19 and Altai*, respectively with a trend towards lower normalised gene diversity in Vindija 33.19). This further suggests that immune gene diversity did not decrease over time. We note that differences in the branch lengths leading to each of the archaic humans reflecting the differences in ages of the specimens, i.e. branch shortening, do not have a substantial effect on our analyses as we do not observe a strong positive correlation between the age of archaic individuals and the normalised gene diversity (Pearson correlation coefficient between normalised mean SNP/FD ratio and age, using Altai*: 0.08). [Approximate position of Figure 2]

### High MHC gene diversity in archaic humans

MHC genes are known to be among the most diverse genes in the genome, due to the action of long-term balancing selection (Bitarello et al. 2018; Key et al. 2014b; Meyer and Thomson 2001). In contrast to the overall gene diversity which is consistently lower in archaic than in present-day individuals, the levels of gene diversity in MHC genes of archaic humans are comparable to the levels observed in present-day humans (Figure S3). To better understand this signature, we evaluated MHC gene diversity for the three archaic humans and the 14 present-day humans by averaging the normalised SNP/FD ratios (Figure 3 (a)). Both archaic and present-day humans show higher diversity in MHC genes than in the background gene set (indicated by log_2_-values of lower 95% confidence intervals > 0). Interestingly, MHC diversity is about 47-fold higher than the background genes in archaic humans (95% CI: 32 to 76-fold) but only approximately 7-fold higher than background genes in present-day humans (95% CI: 5 to 9-fold). This higher diversity in the MHC observed in archaics compared to present-day humans is driven largely by the MHC class II genes (Figure 3 (b, c, d)). It is interesting that the normalised gene diversity in the MHC of the two early modern humans Loschbour (which is approximately 7,000 years old (Lazaridis et al. 2014)) and Ust’-Ishim (which is approximately 45,000 years old (Fu et al. 2014)) is comparable to that of present-day humans (Figure 3 (e)), and thus lower than that of the archaic humans – this is also true for the set of innate immunity genes (Figure S4).

**Fig. 3:**
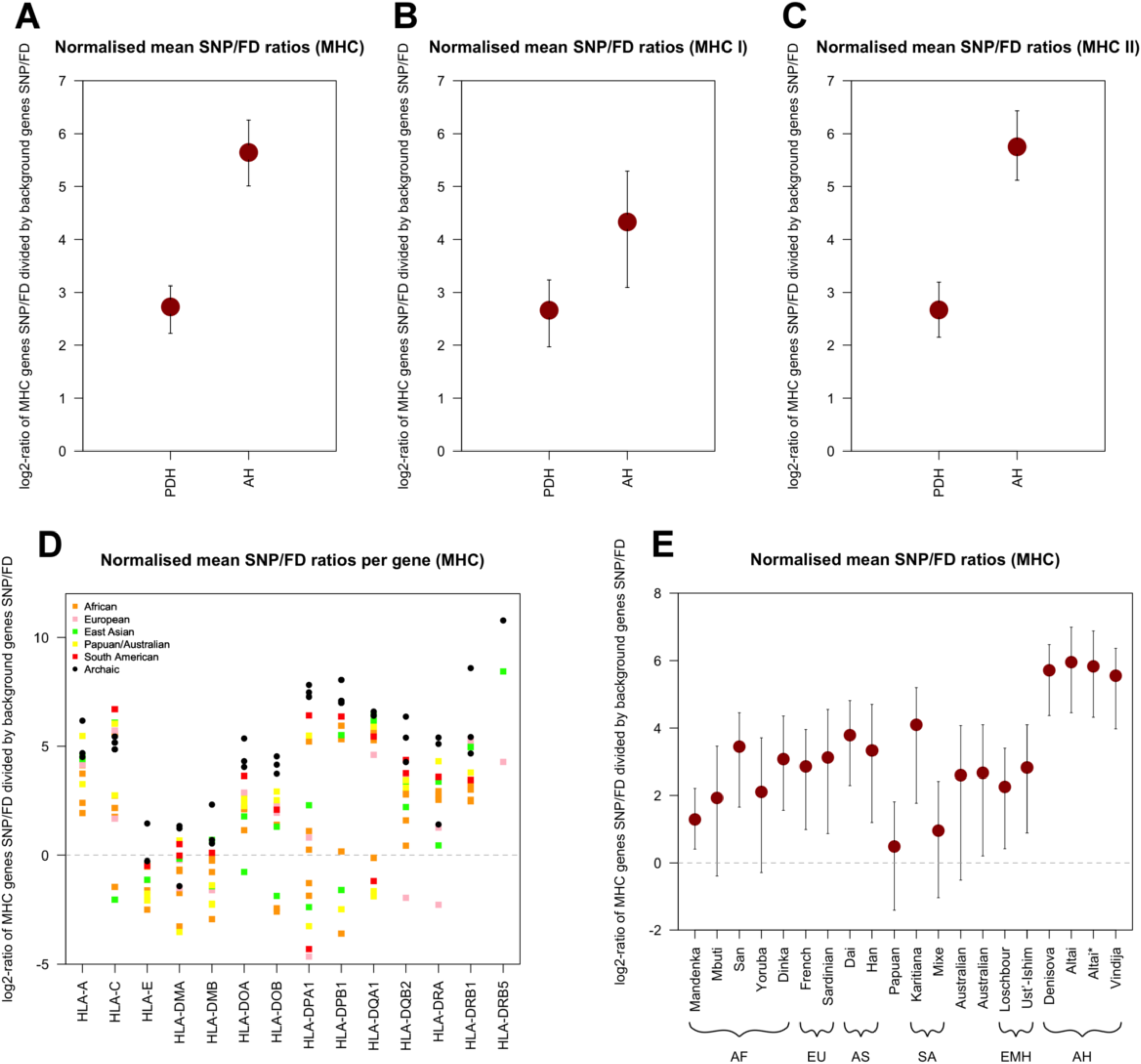
Comparison of normalised mean SNP/FD ratios (log2) of present-day humans (PDH) and archaic humans (AH). **(A)** Average values for PDH and AH for all MHC genes. Error bars give 95% confidence intervals calculated via bootstrap (B = 5,000). **(B)** Average values for PDH and AH for all MHC class I genes. **(C)** Average values for PDH and AH for all MHC class II genes. **(D)** Distribution of SNP/FD ratios (log2) of MHC genes (x-axis) for all individuals. Missing values for single individuals can either be genes without FDs or genes with fewer than 500 callable sites. **(E)** Comparison of normalised mean SNP/FD ratios (mean) between single individuals. AF = Africa, EU = European, AS = Asian, SA = South American, EMH = Early anatomically Modern Humans, AH = Archaic Human. Note differences in the y-axis. Dashed lines in **(D)** and **(E)** give expected values if the mean values for innate immunity and autosomal protein-coding background genes are the same.

Comparing MHC gene diversity between archaic and early modern humans also helps to determine whether problems with the alignment of short ancient DNA reads in these highly polymorphic genomic regions could lead to an over-estimate of diversity. The sequences generated from these ancient specimens have comparable read length distributions (Figure S5 (a)), and a similarly high median genomic coverage to the three archaic genomes (Figure S5 (b)). Since MHC gene diversity in the Loschbour and Ust’-Ishim individuals is no higher than in present-day humans, the high MHC diversity in the archaics is likely not caused by problems with aligning short ancient DNA reads. The fact that the signature is unique of the MHC further suggests that it is not an artefact of incorrect short read mapping of short ancient DNA reads (Figure S6-S8).

The relatively high MHC diversity in archaic humans is evenly distributed in introns and exons, which do not show significantly different SNP/FD ratios (Figure S9). In fact, neither introns nor exons show different SNP/FD ratios between archaic and present-day humans (Figure S10). High gene diversity is thus homogeneously distributed within single genes, rather than due to peaks of diversity in highly polymorphic sections of the genes, which we would expect with mapping errors. The observed patterns are consistent with the high linkage disequilibrium and low average recombination rates of the MHC region (de Bakker et al. 2006; International HapMap 2005; Miretti et al. 2005; Traherne 2008) resulting in high levels of diversity across the entire MHC region. Further, we note that coverage is also evenly distributed across genes – we find no evidence for different coverage in introns than exons – and coverage is also comparable between ancient and present-day individuals (Figure S11).

We separately investigated SNP/FD ratios for the ß-2-microglobulin gene (*B2M*). *B2M* is part of the class I MHC (light-chain) but – unlike the other MHC genes – is located on chromosome 15. It is known to be non-polymorphic in humans (Corazza et al. 2004; Esposito et al. 2008). In concordance with this, we find low SNP/FD ratios that are comparable between archaic and present-day humans (Figure S12). [Approximate position of Figure 3]

### Genes with highest/lowest diversity show similar GO enrichments in archaic and present-day humans

When evaluating the diversity of the entire gene set, genes with the highest diversity (in the top-5% tail of the empirical distribution for SNP/FD ratios) show a highly significant enrichment only of GO categories related to the Major Histocompatibility Complex (MHC) in both the archaic and present-day humans after correcting for multiple testing (Bonferroni correction, k = 17, Table 1). This signal was consistently found when testing the archaic humans in pairs (rather than triplets, Table S1-S3) and individually for the Vindija Neandertal and the Denisova (Table S4-S5), with the Altai Neandertal showing non-significant enrichment (Table S6). [Approximate position of Table 1]

**Table 1:** Significantly enriched GO categories for genes from the top 5% tails of the of SNP/FD empirical distribution in the three archaic humans and three present-day humans (Yoruba, French, Han) ordered by Family-Wise Error Rate (FWER). FWER values are given after correcting for multiple testing (Bonferroni correction, k = 17). Analyses considering pairs of individuals, and single individuals, are presented in the SOM. GO categories related to the MHC region are highlighted with bold font.

|  | Archaic humans (N = 3) |  |  | Present-day humans (N = 3) |  |  |
| --- | --- | --- | --- | --- | --- | --- |
|  | GO ID | GO Name | FWER | GO ID | GO Name | FWER |
| Top 5% | **0032395** | **MHC class II receptor activity** | **0** | 0004984 | Olfactory receptor activity | 0 |
|  | **0042613** | **MHC class II protein complex** | **0** | **0042613** | **MHC class II protein complex** | **0** |
|  | 0004984 | Olfactory receptor activity | 0.017 | **0032395** | **MHC class II receptor activity** | **0.034** |
|  | **0042611** | **MHC protein complex** | **0.034** |  |  |  |

In the bottom-5% tail of the empirical distribution we found enriched categories related to virus-related or mitochondrial functions (Bonferroni correction, k = 17, Table 2). It is tempting to interpret these findings as differences between archaic and present-day humans, especially as previous work suggests that genes related to anti-viral defence are often subject to natural selection, either under strong purifying (Deschamps et al. 2016) or positive selection (Enard et al. 2016; Key et al. 2014a; Manry et al. 2011). However, there was no consistent enrichment pattern when analysing genomes in pairs or individually (rather than triplets), and most enrichments were non-significant trends in some, but not all, archaic or present-day humans (Table S7-S22). Thus, we have no strong evidence of any different GO enrichment patterns between the modern and archaic genomes.

**Table 2:** Significantly enriched GO categories among the bottom 5% tails of the of SNP/FD empirical distribution in the three archaic humans and three present-day humans (Yoruba, French, Han) ordered by Family-Wise Error Rate (FWER). FWER values are given after correcting for multiple testing (Bonferroni correction, k = 17). Analyses considering pairs of individuals, and single individuals, are presented in the SOM.

|  | Archaic humans (N = 3) |  |  | Present-day humans (N = 3) |  |  |
| --- | --- | --- | --- | --- | --- | --- |
|  | GO ID | GO Name | FWER | GO ID | GO Name | FWER |
|  | 0004984 | Olfactory receptor activity | 0 | 0003676 | Nucleic acid binding | 0 |
| **Bottom 5%** | 0004930 | G-protein coupled receptor activity | 0 |  |  |  |
|  | 0004888 | Transmembrane signalling receptor activity | 0 |  |  |  |
|  | 0098800 | Inner mitochondrial membrane protein complex | 0 |  |  |  |
|  | 0099600 | Transmembrane receptor activity | 0 |  |  |  |
|  | 0005125 | Cytokine activity | 0 |  |  |  |
|  | 0005179 | Hormone activity | 0 |  |  |  |
|  | 0098798 | Mitochondrial protein complex | 0 |  |  |  |
|  | 0038023 | Signalling receptor activity | 0 |  |  |  |
|  | 0007186 | G-protein coupled receptor signalling pathway | 0.017 |  |  |  |
|  | 0005743 | Mitochondrial inner membrane | 0.034 |  |  |  |

Together, our results indicate that the enrichment of gene categories among the genes with the highest or lowest diversity is not specific to archaic nor present-day humans, with patterns of diversity in genes with the highest or lowest SNP/FD ratios probably being shaped by the action of long-term balancing selection and strong purifying selection or selective sweeps, respectively. However, we caution that the strength of our conclusions is limited by our small sample size, and that they will have to be confirmed when more archaic genomes become available. [Approximate position of Table 2]

## Discussion

Our results are consistent with previous studies that have reported lower genetic diversity in archaic humans than in present-day humans, both genome-wide and in protein-coding regions (Castellano et al. 2014; Meyer et al. 2012; Prufer et al. 2017; Prufer et al. 2014). In a recent study, lower protein-coding diversity observed in a set of 73 innate immunity genes (Sullivan et al. 2017) was interpreted as suggesting that Neandertals may have lacked the functional immune diversity necessary to survive new pathogen infections (Houldcroft and Underdown 2016; Sullivan et al. 2017; Wolff and Greenwood 2010). Here, we re-evaluated this hypothesis by studying the diversity of a set of 1,548 innate immune genes, and by explicitly comparing them to all autosomal protein-coding genes. Using this set of innate immune genes, we find no significant difference in diversity between protein-coding genes involved in innate immunity and all other autosomal protein-coding genes in any present-day or archaic individual. More strikingly, we see no difference in innate immune gene diversity between the older Altai Neandertal and the younger Vindija Neandertal individuals who lived at least 70,000 years later, as might have been the case if Neandertals were losing important gene diversity over time. A larger number of Neandertal genomes are needed to confirm our results, but with the current available genomes, we find no evidence to link a specific reduction in innate immune gene diversity to Neandertal extinction. We cannot exclude, though, that the global reduction in genome-wide diversity in archaic humans affected the function of immune related genes.

As expected from long-term balancing selection, we find that diversity in MHC genes is much higher than the diversity in other autosomal protein-coding genes (Bitarello et al. 2018; Key et al. 2014b; Meyer and Thomson 2001) but, interestingly, this effect is much stronger in archaic than in present-day humans: for archaic humans, we find an approximately 47-fold higher diversity in MHC than in the protein-coding background, whereas for present-day humans, MHC diversity is only about 7-fold higher than in background genes. This signal is driven by very high diversity in the polymorphic MHC class II genes. This is consistent with the analysis of 12 MHC genes by Sullivan et al. (2017) who reported a significantly higher number of non-synonymous SNPs in Neandertals compared to present-day humans, also at intermediate frequencies, for MHC genes relative to a genome-wide background. From this, they concluded that heterozygote advantage at MHC loci might have been stronger than expected and might have maintained crucial functional variation despite low N_e_ in Neandertals (Sullivan et al. 2017). Interestingly, the two early modern humans we analysed here show similar MHC gene diversity to present-day people (Figure 3 (e)), even though their population densities, and therefore likelihoods of pathogen transmission, were presumably more similar to that of archaic humans than to that of present-day humans. This is not completely unexpected as the effective population size of early modern humans was likely higher than that of archaic humans (Fu et al. 2014). However, it contrasts with the unexpectedly high MHC diversity maintained in the archaic genomes.

Although it is difficult to completely rule out that technical artefacts might increase our measure of diversity in the MHC genes, none of our tests show evidence for mis-alignments of short ancient DNA reads being responsible for our findings. There are thus two plausible explanations for the pattern of MHC diversity. (i) It could be caused by the old TMRCA in the MHC region (Leffler et al. 2013; Tesicky and Vinkler 2015) and the persistent presence of intermediate frequency alleles as a consequence of long-term balancing selection (Bitarello et al. 2018; Key et al. 2014b; Meyer and Thomson 2001) resulting in the maintenance of sequence diversity in these genes which has been reported for targets of long-term balancing selection (Bitarello et al. 2018). (ii) It could have been shaped by stronger selective pressures in Neandertal than humans preventing extensive loss of diversity in these genes – although in populations with small N_e_ the effects of selection are generally weaker than in population of larger N_e_ (Charlesworth 2009; Hoffmann et al. 2017; Willi et al. 2006). A possible mechanism for this would be associative overdominance. In that case, selection against homozygous recessive deleterious alleles in genomic regions could result in overdominance at linked neutral loci, boosting the effects of balancing selection. Associative dominance has recently been reported to drive maintenance of genetic diversity in experimental small N_e_ populations of field-caught *Drosophila melanogaster*, especially in regions with low recombination rates (Fraser 2017; Schou et al. 2017). This is particular interesting as recombination rates in the human MHC region on average are notably lower than expected from the genome average (de Bakker et al. 2006; International HapMap 2005; Miretti et al. 2005; Traherne 2008). Theoretically, the increased diversity in the MHC could also be the result of introgression into the archaic hominins. However, we note that (i) gene flow of this magnitude has not been detected to date and (ii) if introgression contributed, we would not expect it to strongly affect the gene set as a whole. Therefore, we consider this an unlikely explanation.

Future sequencing of additional high-coverage archaic genomes that sample the geographic and temporal distribution of Neandertals will allow questions about the effects of gene diversity on Neandertal fitness to be addressed in greater detail.

## Authors’ Contributions

AMA and JK designed the study. DR analysed the data. All authors interpreted results. DR wrote the manuscript with contributions from FMK, AMA and JK. All authors read and approved the final manuscript.

## Acknowledgments

We thank Steffi Grote for help with the GO enrichment analysis, and Bárbara D Bitarello, Michael Dannemann, Steffi Grote, Benjamin M Peter, Svante Päabo and Joshua M Schmidt for helpful comments. We are also grateful for comments of two anonymous reviewers. This work was supported by the Max Planck Society.

## Supplementary Material

Supplementary data is available online.

